# Stress-Induced *PTBP1* Reprograms Neuronal Function and Activates Cellular Senescence

**DOI:** 10.64898/2026.04.03.711742

**Authors:** Priyanka Priyanka, Amir Gamliel, Havilah Taylor, Kenneth Ohgi, Michael G. Rosenfeld

**Affiliations:** School and Department of Medicine, University of California San Diego, La Jolla, CA 92093, USA; Cellular and Molecular Medicine, Department of Medicine, University of California San Diego, La Jolla, CA 92093, USA

## Abstract

Chronic oxidative stress is a major contributor to neuronal aging. Due to the lack of homologous recombination (HR) DNA damage repair, high oxygen consumption in neurons causes DNA damage accumulation with age, resulting in a decline in neuronal function, senescence-like phenotypes and onset of neurodegenerative diseases. Here, we identify increased *PTBP1* as a stress-inducible negative regulator of neuronal gene expression and senescence-protectant genes. Oxidative stress robustly increases *PTBP1* expression in ShSY-5Y differentiated neurons and primary mouse cortical neurons, coinciding with the loss of neuronal genes, including neuronal *PTBP2*, and activation of stress-responsive genes. Knockdown of *PTBP1* in fibroblasts reduces the expression of key senescence genes. Transcriptomic analyses revealed that *PTBP1* overexpression results in coordinated shift in gene expression characterized by repression of neuronal commitment genes and activation of stress and senescence genes. Mechanistically, *PTBP1* induction is regulated by stress induced CTCF binding at the *PTBP1* promoter. Together, our findings suggest that alteration in levels of *PTBP1* acts as a molecular switch between neuronal function and survival, providing insight into transcriptional adaptations associated with aging.

**SUMMARY:**

- Loss of *PTBP1* in fibroblasts acts as a senescence protective gene
- xidative stress induces expression of *PTBP1*, reducing neuronal function gene expression and activating stress and cell cycle genes
- Ectopic *PTBP1* expression reprograms neuronal transcription, down-regulating cell fate commitment genes and activating a cell senescence program
- xidative stress induces *PTBP1* and suppresses neuronal specific *PTBP2* expression in primary cortical neurons

## INTRODUCTION

Ageing is characterised by a progressive accumulation of oxidative damage, decline in cellular function, and a loss of homeostasis [1–4]. Neurons are constantly under oxidative stress due to high metabolic oxygen consumption and psychological factors, that contribute to impaired neuronal function and vulnerability to degeneration [2–4,6]. While these stress responses are essential for survival and effective functioning, they involve trade-offs that favour survival at the expense of cellular differentiation, particularly in post-mitotic neurons (López-Otin et al., 2013; Mattson & Arumugam, 2018). Aged and diseased neurons undergo degradation of identity, loss of function, and eventually die, leading to cognitive decline with age and in neurodegenerative diseases. Memory formation involves synaptic activity, which is associated with higher oxygen consumption, metabolic demands and oxidative damage. However, with age, neurons accumulate DNA damage due to a decline in DNA repair efficiency and turnover mechanisms [1–4]. Functional impairment of cortical pyramidal neurons of mice leads to dendritic and synaptic degeneration, resulting in deficits in learning, memory, and sensory integration, and increased network hyperexcitability, that is associated with various neurodegenerative conditions, including Alzheimer’s disease and Frontotemporal dementia (FTD).

Several RNA binding proteins (RBPs) play critical role in neuronal differentiation and maintenance, including HuD, nELAVs, RBFOX1, and PTBP1, by controlling mRNA splicing, stabilizing, transporting, and locally translating mRNA in axons and dendrites, thereby helping sustain neuronal identity and synaptic function [8–11]. By rapidly adjusting the translation of repair and pro-survival mRNAs at sites of stress or injury, RBPs support neuronal resilience and contribute to regeneration-associated responses in both development and disease contexts [8–11]. Recently, *PTBP1* (polypyrimidine tract-binding protein 1) has emerged as a pivotal RNA-binding regulator [9–11]. During neuronal maturation, *PTBP1* expression is downregulated, and its paralog *PTBP2* increases, enabling neuronal-specific splicing programs. Early studies demonstrated that ASO-mediated *PTBP1* depletion in the midbrain converts astrocytes into dopaminergic neurons and rescues Parkinson’s disease phenotypes by enabling axonal regeneration to reconstruct the nigrostriatal circuit, elevating *PTBP1* as a putative “gatekeeper” of neuronal fate [12–16]. However, potential reprogramming from astrocytes to neurons remains controversial and requires additional scientific evidence (Qian, Kang et al. 2020).

Several studies have reported elevated *PTBP1* and decreased *PTBP2* expression in aged human brains, suggesting that this imbalance may contribute to stress-related RNA processing abnormalities and neuronal vulnerability in the pathophysiology of aging and neurodegeneration [9–11]. With age, neurons accumulate both damage and *PTBP1* expression with age, neurons accumulate both damage and PTBP1 expression [1–4,17]. However, how *PTBP1* expression is regulated in the context of cellular stress, age and how this impacts neuronal function, remains unclear

Here, we investigate the role of *PTBP1* in mediating neuronal responses to oxidative stress and neuronal function. We show that oxidative stress robustly induces *PTBP1* expression in both neuronal cell models and mouse primary cortical neurons. Through transcriptomic and functional analyses, we demonstrate that *PTBP1* overexpression drives a coordinated transcriptional shift characterised by repression of neuronal function-associated genes and activation of cell proliferation and cellular senescence pathways. Furthermore, we provide evidence that this stress-induced upregulation of *PTBP1* is mechanistically regulated by increased CTCF binding at its promoter. Importantly, depletion of *PTBP2* does not recapitulate these effects, highlighting a specific role for *PTBP1*. Together, our findings identify *PTBP1* as a stress-responsive regulator that promotes a shift toward a stress-adaptive, ageing-like transcriptional state in neurons.

## RESULTS

### *PTBP1* repression acts as a senescence-protectant gene in human fibroblasts

Human IMR-90 Fibroblast cells undergo chemical-induced senescence by treating cells with 250nM doxorubicin for 24h, followed by 72h of wash off to induce Senescence, in either control shRNA or *PTBP1* knockdown cells. *PTBP1* knockdown results in reduced expression of key senescence-associated genes *Cdkn1a (*P21*), Cdkn2a* (p16)*, MMP3, NFAT5, IL-6* and reduced β-gal staining (**Fig. 1A, 1B, 1C**). These results suggest that *PTBP1* inhibition can protect the cells from induced senescence, as indicated by β-gal staining. These results suggest that PTBP1 expression plays a role in cellular senescence, and its diminished expression in postmitotic neurons might serve a protective mechanism.

**Figure 1.**
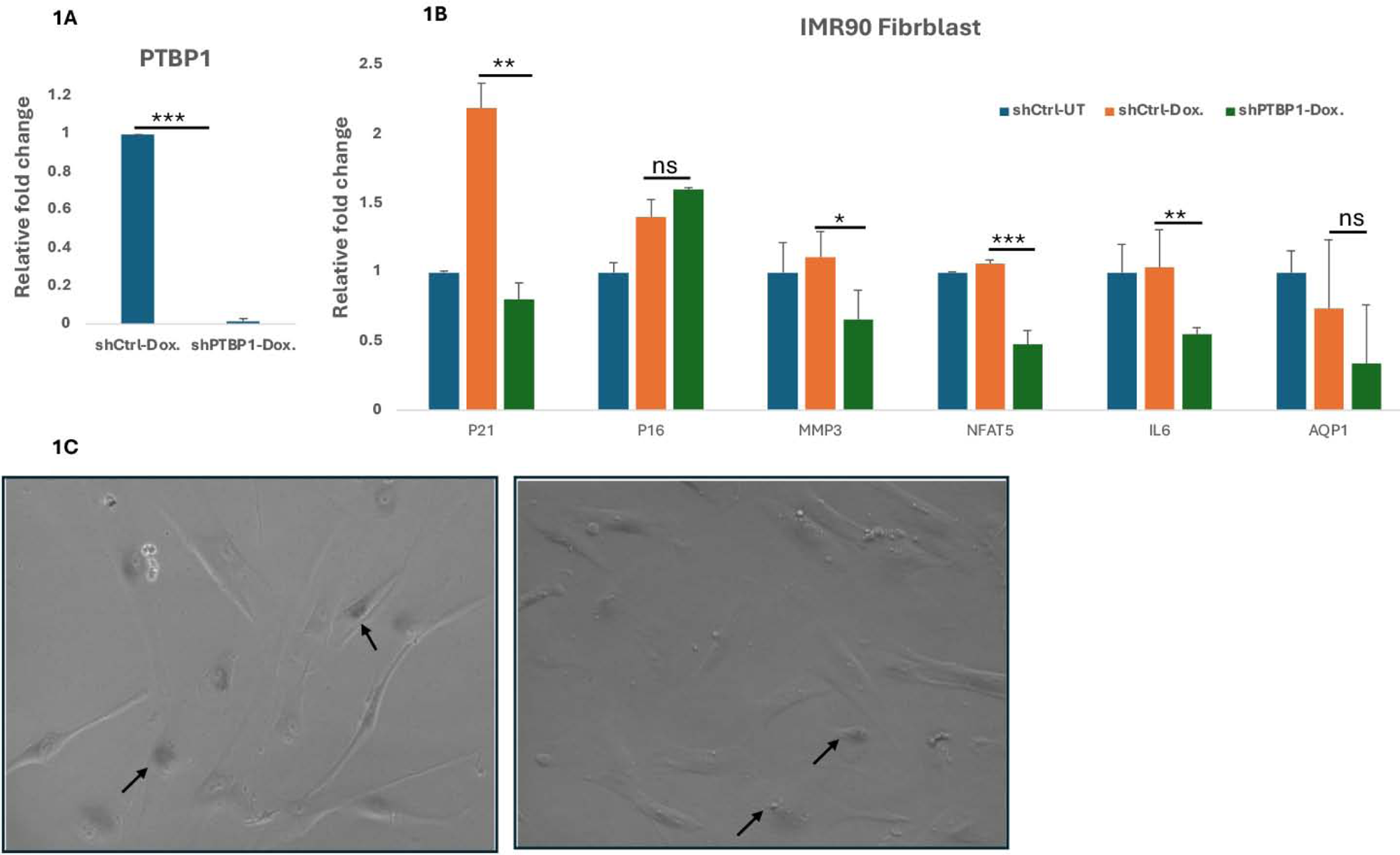
PTBP1 Knockdown in fibroblasts acts a Senescence Protectant gene. (A–B) Relative expression of genes by qPCR in IMR90 human fibroblasts, either left untreated or exposed to 250 nM doxorubicin for 24 h, washed out 72 h, followed by RNA isolation and β-Senescence-Associated galactosidase (SA-gal) staining. Gene expression levels were quantified by RT–qPCR and normalised to control conditions. PTBP1 knockdown attenuates stress-induced transcriptional changes associated with senescence. (C) β-Gal staining of senescence-associated markers under control and PTBP1 knockdown conditions following doxorubicin treatment, demonstrating a protective effect of PTBP1 depletion against stress-induced senescence. Data are presented as mean ± SEM from biological replicates. Statistical significance was determined using Student’s t-test with *p < 0.05, **p < 0.01, ***p < 0.001, ns = not significant.

### Oxidative stress induces *PTBP1* expression in SH-SY5Y differentiated neurons

Therefore, to understand the relevance of *PTBP1* suppression in neurons, we used SH-SY5Y neuroblastoma cells that were differentiated into neurons by initial 4-day treatment with 10 µM of Retinoic Acid (RA) followed by 5 days of 50 ng/mL Brain-Derived Neurotrophic Factor (BDNF) to achieve *in vitro* differentiated dopaminergic-like post-mitotic neurons (DIV) (**Fig. 2A**). qRT-PCR revealed the increased expression of neuronal genes including- *TUBB3, NRN1, RBFox3, ENO2, MAPT* confirms the expression of neuronal markers. Western blot and qPCR confirmed significant reduction in PTBP1 protein & mRNA expression, respectively (**Fig. 2B**, **2C)**. These neurons were exposed to oxidative damage by treating with 0.25 mM H_2_O_2_ for 15-minute pulses, followed by wash off. qRT-PCR showed an significant increase in the *PTBP1*, *Cdkn1a*, and *REST* expression, and decreased levels of *MAP2* and neuronal genes (**Fig. 2D**, **2E)**. RE1-Silencing Transcription repressor (REST) is the key transcription repressor that suppresses expression of neuronal genes, and its expression is very low in mature neurons, similar to *PTBP1*. Similarly, elevated *Cdkn1a* expression upon H_2_0_2_. Interestingly, *REST*, *CDKN1A*, and *PTBP1* dysregulated expression was reported in Alzheimer’s and Parkinson’s diseases and in aged human neurons (Lu et al., 2014, *Nature*; Morris et al., 2008, *J Neurosci*; Qian et al., 2020, *Nature*).

**Figure 2.**
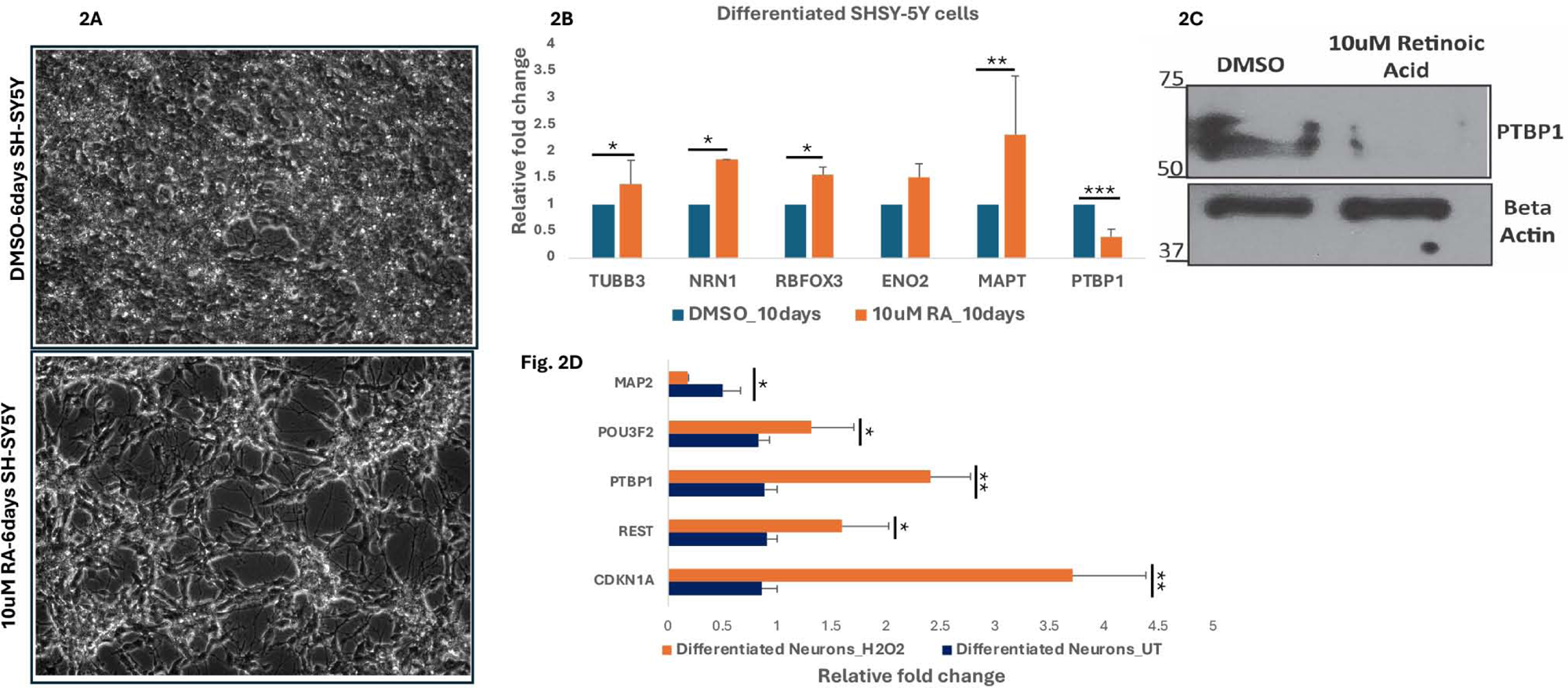
Oxidative stress induces *PTBP1* expression in SH-SY5Y differentiated neurons. (A) Human neuroblastoma SH-SY5Y cell lines were incubated with either DMSO or (undifferentiated control) or 10 μM retinoic acid (RA) for 4-6 days, showing differentiation and neurite formation. (B–C) Relative fold expression of key neuronal genes, including *PTBP1,* was measured using qRT-PCR and protein expression was measured using western blot (C) in undifferentiated and RA-differentiated cells. (D) Relative expression analysis of neuronal function–associated genes under oxidative stress conditions, demonstrating their downregulation concomitant with *PTBP1* induction. Data represent mean ± SEM from independent experiments. Statistical significance was assessed using the Student’s t-test.

### Oxidative stress results in loss of neuronal function genes and activates stress and cell proliferation genes

Cognitive and motor decline with age is linked to Cellular Senescence (SASP) accumulation, accumulated damage in the prefrontal cortex and in cortical neurons. Therefore, we isolated primary mouse cortical neurons and cultured them *in vitro* media for around 5 days to understand the relevance of oxidative stress-induced *PTBP1* genes (**Fig. 3A**). We exposed mouse primary cortical neurons to 0.25mM H_2_O_2_ and performed RNA sequencing which revealed extensive transcriptional reprogramming. We found 755 (3.005%) genes had significantly reduced expression, including the key neuronal function genes such as *PTBP2, MAP2, Rbfox3, ENO2, and BDNF* (**Fig. 3E**). Gene ontology (GO) term analyses confirmed enrichment of genes involved in chemical synaptic transmission, neuroactive ligand-receptor interaction, and chemical synaptic transmission among the most significantly downregulated pathways (**Fig. 3D**). While transcriptomic analyses revealed that 1942 (7.730%) of the Genes were significantly upregulated (log2 fold change> 1, FDR < 0.05), including *PTBP1*, *REST*, *Cdkn2b*, *Rad51b*, *CdK6*, *CCNA2*, and several genes are involved in DNA damage repair and cell cycle (**Fig. 3E**). GO term analyses show that the pathways involved in cell population proliferation, cellular response to stimuli, and response to wounding were significantly affected (**Sup Fig. 1B, 1C**). Gene ontology (GO) analysis confirmed enrichment of neuronal identity and function genes among downregulated genes, consistent with an increase in stress-associated and cell cycle proliferation genes that reflects a shift away from the differentiated neuronal state. We confirm the expression of key genes using qRT-PCR, showing stress-induced increase in *Cdkn1a, PTBP1, and REST* expression in mouse primary cortical neurons (**Fig. 3B**, **Sup Fig 1A**). Interestingly, neuronal-specific *PTBP2* expression was significantly reduced by qRT-PCR (**Fig. 3C**).

**Figure 3.**
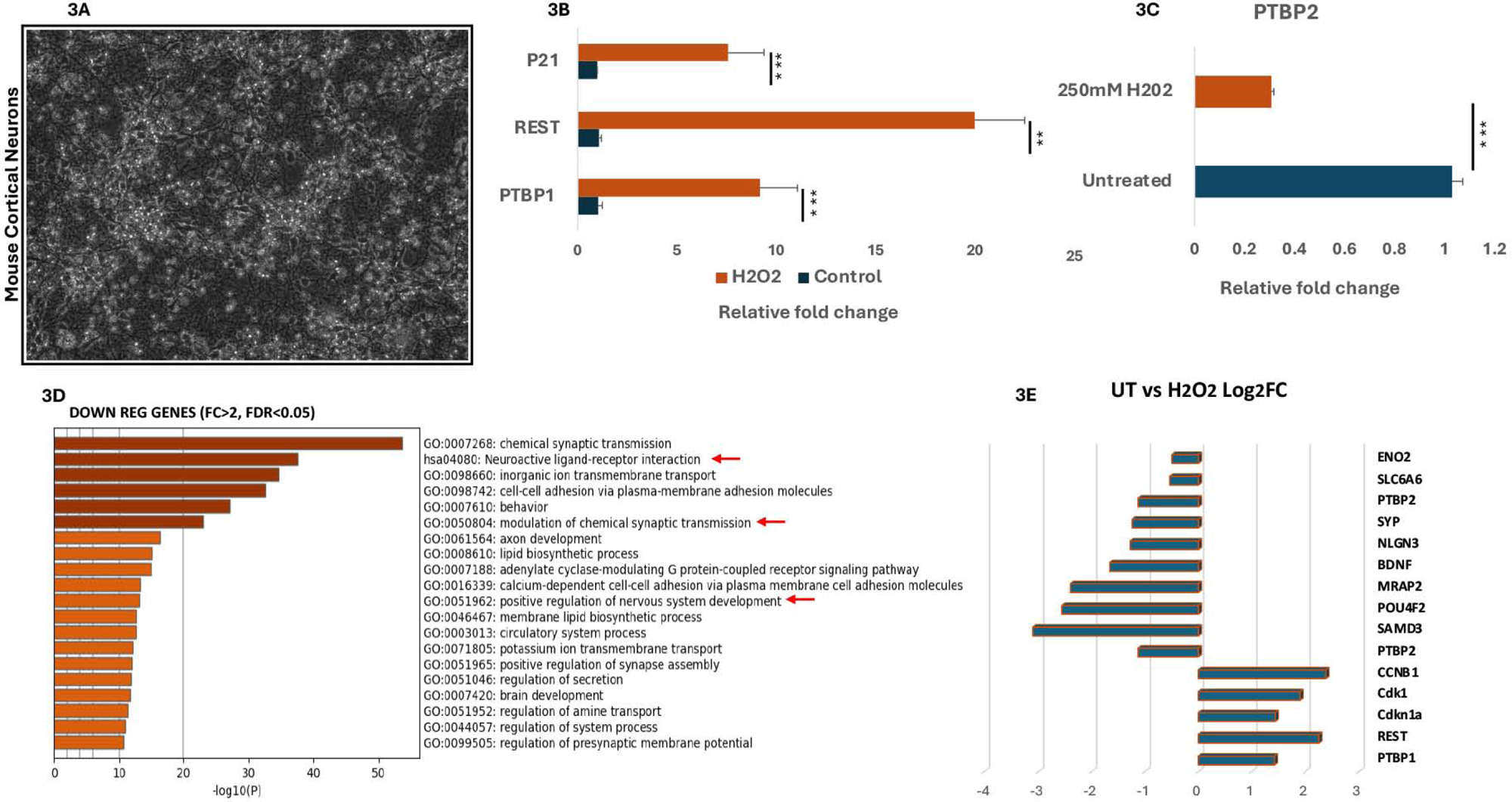
Oxidative stress induces PTBP1 and suppresses neuronal function gene expression in primary cortical neurons. (A) Mouse primary cortical neurons isolated from E15.5 mouse embryo cortex region and grown in a plate for DIV 5. (B) Relative expression of PTBP1 in primary mouse cortical neurons under basal and oxidative stress conditions. (B-C) Relative expression analysis of neuronal function–associated genes, showing significant downregulation following oxidative stress using RT-PCR. (D) Transcriptomic profiling identifying significantly downregulated genes (fold change >2, FDR < 0.05) under oxidative stress conditions. Functional categorisation of downregulated genes highlighting enrichment of neuronal function–related pathways using Metascape. (E) Log2 Fold change expression of a set of neuronal function-related genes, stress and cell proliferation genes. All genes plotted are FDR < 0.05.

### PTBP1 overexpression partially recapitulates stress-induced transcriptional programs

To assess whether stress-induced *PTBP1* expression is causal and sufficient to drive stress-associated transcriptional changes in mouse primary cortical neurons. We ectopically overexpressed *PTBP1* in neuronal cells under basal conditions and in oxidative stress (**Fig. 4A**). we performed the RNA-seq transcriptomic analysis that demonstrated that *PTBP1* overexpression alone induced a subset of genes normally activated by oxidative stress (**Fig. 4B-4C**), including *REST*, *CDKN2A* and 590 other genes. GO term analysis reveals that genes belong to the response to hypoxia, stress and negative regulation of neuronal apoptosis pathways (**Fig. 4D, 4E**). Conversely, *PTBP1* overexpression preconditions cells toward a less differentiated and more stress-adapted transcriptional state.

**Figure 4.**
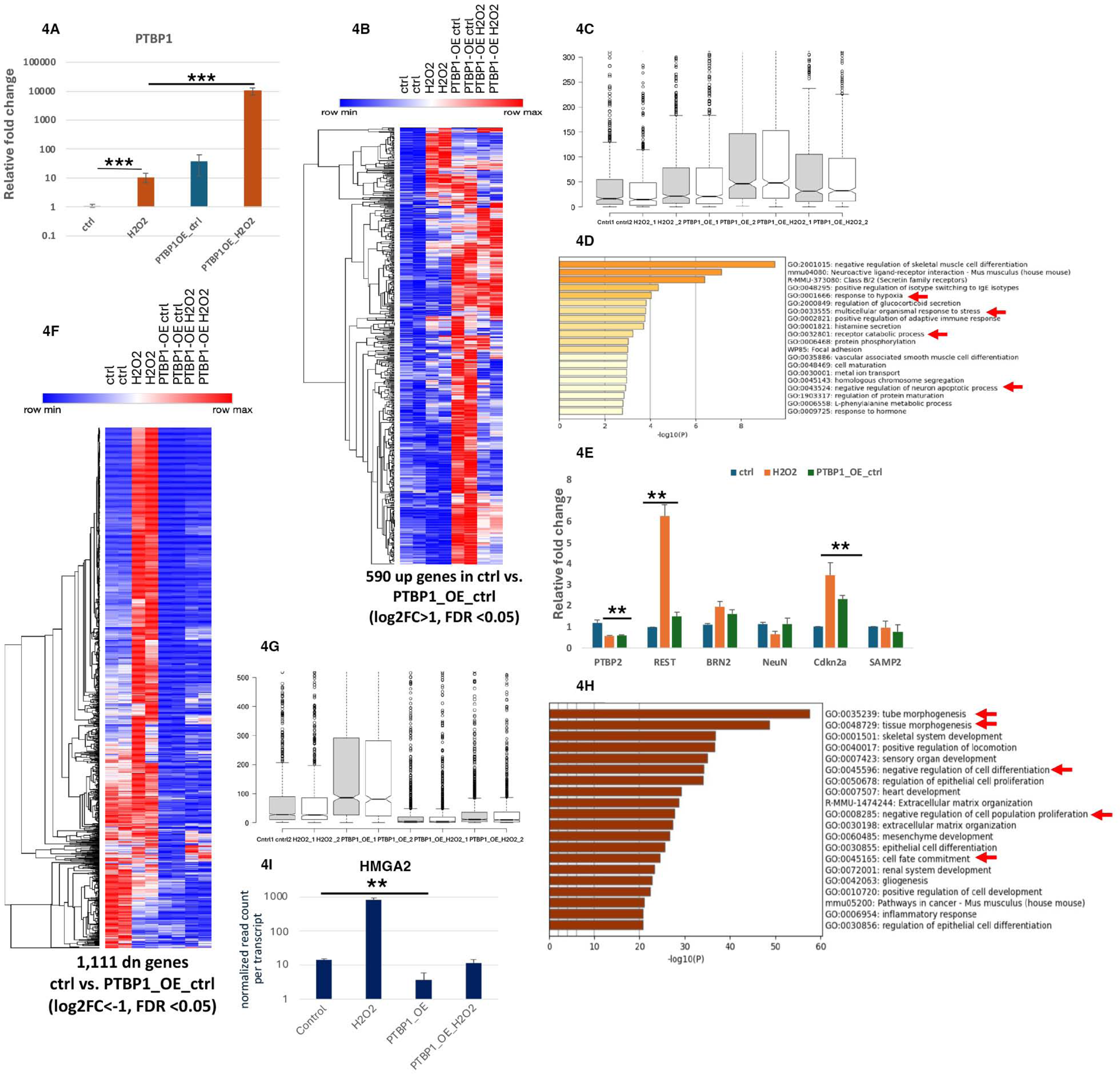
PTBP1 overexpression partially recapitulates stress-induced transcriptional programs. (A) Relative fold change expression of PTBP1 in mouse cortical neurons in either control or in ectopically overexpressed PTBP1 using the PTBP1-Myc tagged vector from Addgene. (B-D) Heatmap of 590 upregulated genes in control vs PTBP1 OE control cells showing various clusters, showing that only PTBP1 OE affects the expression of these genes. Differentially expressed genes in control versus PTBP1 overexpression (OE) conditions, identifying 590 upregulated genes and GO term analysis revealed belong to response to stress, neuronal apoptotic process (log2FC > 1, FDR < 0.05). (E) Validation of selected upregulated genes by RT–qPCR, demonstrating activation of *CDKN2A*, *REST* and suppressed *PTBP2* expression upon *PTBP1* overexpression. (F-I) Heatmap representation of normalized transcript counts showing global expression patterns across conditions: control, oxidative stress, PTBP1 OE, and combined PTBP1 OE with oxidative stress. Transcriptomic analysis identifying 1,111 downregulated genes (log2FC < −1, FDR < 0.05) upon PTBP1 overexpression. Heatmap analysis indicates that these genes remain suppressed under PTBP1 OE and fail to be induced upon oxidative stress, in contrast to their activation in control cells. GO term analysis shows that these genes regulate cell fate commitments, negative regulators of cellular differentiation as shown normalized expression HMG2A gene from RNA-seq data. Collectively, these data demonstrate that PTBP1 overexpression induces a transcriptional shift characterized by repression of neuronal function genes and activation of stress-adaptive programs.

Transcriptomic profiling revealed that *PTBP1* overexpression results in the downregulation of 1,111 genes compared to control cells. Notably, heatmap analysis shows that these genes remain suppressed and are not further induced by oxidative stress, suggesting that *PTBP1* overexpression overrides their normal stress-responsive activation (**Fig. 4F, 4G**). Go term analysis revealed pathways affected are cell fate commitment, negative regulation of cell proliferation and differentiation (**Fig. 4H**). Interestingly, neuronal-specific *PTBP2* expression was significantly repressed as observed upon oxidative stress (**Fig. 4E**). Interestingly, High-mobility Group AT-hook (HMGA2), which represses premature differentiation and promotes self-renewal and neural cell fate by its chromatin condensation activity (Kuwayama, Kujirai et al. 2023), was significantly suppressed upon *PTBP1* overexpression (**Fig. 4I**). Further analysis revealed that around 815 genes were no longer induced upon H_2_O_2_ in *PTBP1* overexpression cells, suggesting that elevated *PTBP1* expression drastically changes neuronal transcriptomics (Sup Fig. 2A). We plotted 1700 genes expression involved in neuron generation from Mouse Genome Informatics portal revealing that *PTBP1* overexpression drastically dysregulates the expression of neuronal genes (**Sup Fig. 2B**).

### *PTBP2* knockdown does not phenocopy the *PTBP1* Overexpression

Based on the observation that both oxidative stress and *PTBP1* overexpression results in suppression of *PTBP2* in mouse cortical neurons. To check whether the shift from neuronal differentiation to neuronal stress and a decrease in cell fate commitment genes might be the result of reduced *PTBP2*, we knocked down the *PTBP2* expression using siRNA to determine whether it does affect *PTBP1* basal or stress-induced expression (**Fig. 5A**). RNA-seq analysis of *PTBP2* knockdown cells does not affect the neuronal function genes, but cellular repair genes were repressed (**Sup Fig. 2C, 2D, 2E**). These data suggest that *PTBP2* knockdown did not recapitulate *PTBP1* overexpression and effects on neuronal function genes.

**Figure 5.**
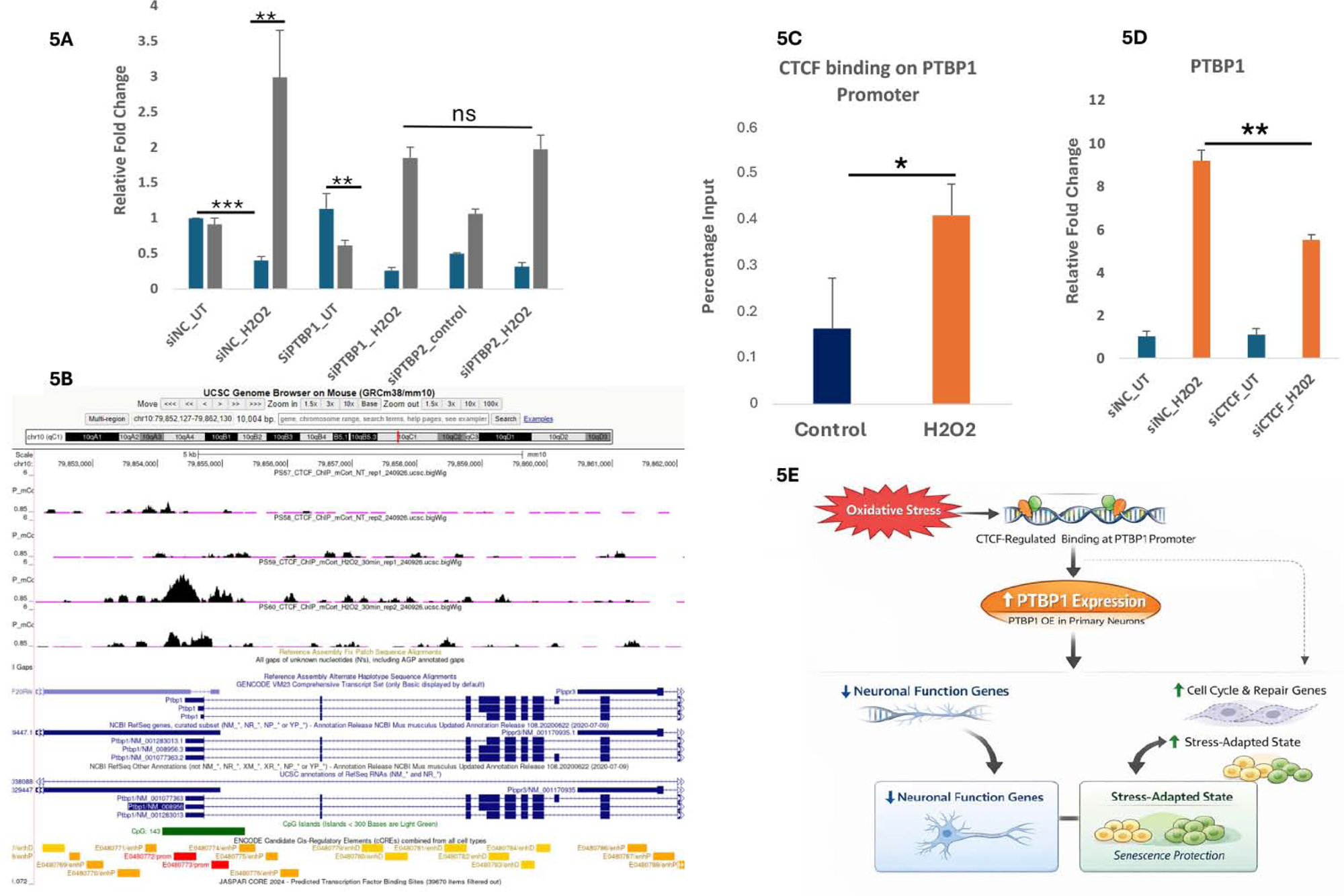
CTCF Chromatin binding regulates stress-responsive PTBP1 gene activation. (A) Relative Fold change expression in mouse cortical neurons either transfected with control siRNA or siRNA against PTBP1 or PTBP2, and relative fold change expression of PTBP1 (grey) and PTBP2 (blue) was measured in either UT or treated with 250nM H_2_O_2_ for 15 mins. (C) UCSC screenshot of CTCF occupancy at the PTBP1 locus under control and oxidative stress conditions using CTCF ChIP-seq. (D) ChIP-qPCR showing increased CTCF binding on the PTBP1 locus in mouse cortical neurons. (E) CTCF perturbation using siRNA in either UT or in H_2_O_2_ *PTBP1* expression was measured using qPCR. Relative fold change of PTBP1 expression, CTCF partially regulates its stress-induced upregulation. Data are presented as mean ± SEM. Statistical analysis was performed using Student’s t-test.

### CTCF Chromatin binding regulates stress-responsive *PTBP1* gene activation

To understand the mechanism behind the oxidative stress-induced *PTBP1* expression, we investigated regulation of *PTBP1* promoter locus upon oxidative stress [18–21]. We identified stress-induced binding of the chromatin architectural protein CTCF on the *PTBP1* promoter in mouse primary cortical neurons using CTCF ChIP-Seq, which was confirmed by ChIP-qPCR (**Fig. 5B, 5C**). siRNA-mediated knockdown of CTCF expression altered stress-induced *PTBP1* expression, suggesting that chromatin organisation contributes to stress-responsive regulation of *PTBP1* expression (**Fig. 5D**).

## DISCUSSION

Our study identifies *PTBP1* expression induced in response to oxidative stress, which promotes a transcriptional program consistent with a stress-adaptive but functionally compromised state. Stress-induced *PTBP1* expression drives the coordinated transcriptional shift from differentiated or lineage-specific genes to activation of repair and cellular senescence pathways, reflecting the fundamental homeostatic erosion associated with ageing and neurodegenerative diseases [1–4,6,7,18–21]. *PTBP1* knockdown reduces the doxorubicin-induced senescence gene markers and β-gal staining in human fibroblast cells, reflecting that its suppression protects cells from induced senescence, even in dividing cells.

Interestingly, exposure to oxidative stress robustly induces *PTBP1* expression in both ShSY-5Y differentiated human neuronal cells and in mouse primary cortical neurons. Notably, *PTBP1* expression is suppressed during neuronal differentiation, suggesting that its re-expression under stress reflects a shift toward a less differentiated and more inflammatory cellular state, consistent with our transcriptomic and functional analyses (**Fig. 5H**) [8–11,25]. Such reprogramming is characteristic of aging and stress-associated conditions, where the preservation of specialized cellular identity is compromised in favour of survival [1–4,7, 22–24]. In this context, *PTBP1* induction may act as a conserved stress-adaptive mechanism in neurons to cope with oxidative damage, albeit at the cost of functional integrity (**Fig. 5H**).

Our transcriptomic analyses further reveal that ectopic *PTBP1* overexpression represses genes associated with cell fate commitment, resulting in negative regulation of cellular differentiation, and activates genes involved in repair and stress or senescence pathways. This transcriptomic reprogramming is striking for the post-mitotic state of neurons and is consistent with the abrupt activation of the cell cycle in neurodegenerative disease [26–28] (Herrup & Yang, 2007). Oxidatively-stressed neurons activate repair genes to maintain genomic stability and protect themselves from apoptosis [1–4,7]. These findings suggest that *PTBP1* functions as a coordinated molecular rheostat that balances between maintenance of neuronal phenotype and activation of stress programs, including senescence, reflecting an interesting biological phenomenon of cells choosing to repair and replenish over risking their fate or functions [1–4,18,19,25].

Importantly, *PTBP2* depletion does not phenocopy the effects of *PTBP1* overexpression, underscoring the specificity of *PTBP1* in mediating stress response [9–11,29,30]. Although *PTBP1* and *PTBP2* share structural and functional similarities, their distinct expression patterns and target specificities have been well documented (Boutz et al., 2007; Makeyev et al., 2007) [9–11,29,30]. Our findings extend this distinction by highlighting a role for *PTBP1* in orchestrating stress-induced transcriptional reprogramming in neurons [9–11,18,19,25].

Mechanistically, oxidative stress induces CTCF binding at the *PTBP1* promoter, and CTCF knockdown partially suppresses stress-induced *PTBP1* expression. CTCF is a key architectural protein involved in chromatin organisation and gene regulation and has been implicated in dynamic transcriptional responses to environmental stimuli (Ong & Corces, 2014) [31–35]. Our CTCF ChIP-seq data show increased CTCF binding on the *PTBP1* promoter, facilitating stress-induced *PTBP1* expression, thereby linking genome organisation to transcription and RNA regulatory networks [31–35]. However, CTCF may only partially account for *PTBP1* induction, reflecting the technical limitations of achieving only ∼50% reduction in CTCF mRNA due to poor transfection efficiency in primary cortical neurons.

High-throughput transcriptomic analysis of the aged human brain reports increased *PTBP1* and suppressed *PTBP2* expression (Mazin, Xiong et al. 2013), which might reflect accumulated damage with age. This study provides insight into how stress-responsive pathways can reshape neuronal transcriptional programs in ways that mirror ageing-associated changes. The observed reduction in neuronal function gene expression, coupled with increased activation of cellular senescence, is consistent with the potential presence of neurons with an ambiguous cell phenotype associated with aging, susceptible to neurodegenerative disease [1–4,7,18,19,36].

In conclusion, we identify *PTBP1* as a stress-induced regulator that promotes a shift toward a repair-oriented, aging-like transcriptional state in neurons. By linking oxidative stress, chromatin regulation, and RNA-binding protein function, our work provides a framework for understanding how adaptive responses to stress contribute to the progressive loss of neuronal function observed during aging.

### Limitations of the Study

This study primarily relies on transcriptomic and RNA processing analyses in cell-based systems. Future studies will be required to identify direct *PTBP1* RNA targets and to assess the functional consequences of *PTBP1* activation *in vivo*. Additionally, while chromatin regulation appears to contribute to stress-induced *PTBP1* expression, the precise molecular mechanisms underlying this regulation remain to be elucidated.

## MATERIALS AND METHODS

### Cell Culture, transfections and treatment

The protocol for cell culture followed previous work(Oh, Shao et al. 2021). In brief, we originally purchased IMR-90 and ShSY-5Y, HEK293T cells from ATCC, which were maintained in DMEM (Gibco, 10566) low glucose for IMR-90 supplemented with 10% FBS (FB-11, Omega Scientific) in a 5% CO2 humidified incubator at 37 °C. Cells were examined for mycoplasma contamination every 6-12 months. Lentivirus targeting scrambled control or PTBP1 was collected from transfected HEK293T cells, and IMR-90 cells were infected along with polybrene. 48 hours after infection, cells were selected in medium containing 1µg/ml of puromycin. Stable cells were then exposed to 250 nM doxorubicin for 24 hours and then washed off for 72 hours before collecting for RNA or for β-galactosidase staining as per the kit instructions (CST 9860).

ShSY-5Y cell lines grown in 1% FBS solution along with 10µM concentration of Retinoic acid for or DMSO for 6 consecutive days and changing the medium every other day. After 6 days, cells were grown in phenol red-free neurobasal media (Gibco 12348017) supplemented with 50ng/ml of BDNF for the next 4-5 days until they developed neurite growth. Following it, cells are either left untreated or exposed to 250nM of H_2_O_2_ (Sigma, H1009) for 15 minutes and harvested for RNA Isolation.

### Mouse cortical neurons

Sacrifice mouse (preg E15.5) with CO2, spray belly with 70% EtOH. Remove pups by cesarean surgery, and each embryo’s brain was dissected to collect cortex, excluding ganglionic eminences, olfactory bulbs, and meninges to leave mostly cerebral cortex in ice-cold HBSSG. The cortex is digested with trypsin and DNase I at a 37 °C incubator and then plated on POLY-D-LYSINE pre-coated 6- well plate in Neurobasal medium (Invitrogen 21103-049) supplemented with 1x B27 supplement (50X, Invitrogen 17504044), 1x GLUTAMAX (100X) and 1x gentamicin (1000X). After two days, cells were treated with the final concentration of 2.8 µM Arca to kill the dividing cells. The experiment was performed after 6 days in vitro (6DIV). Transfections of HEK293 cells with siRNA, ASO and plasmids at 60–80% confluency was performed using Lipofectamine 2000 (Life Technologies) following the manufacturer’s instructions. The control si-, sh-, and ASO oligos used in this study include Santa Cruz negative control siRNA (sc-36869), IDT control B ASO (QTE-240580), Sigma Mission siRNA universal control #2 (SIC002).

For inducing oxidative stress, cells were treated with 0.25mM H_2_O_2_ (Sigma, H1009) for 15 mins. After treatment, the H_2_O_2_ containing medium was removed/washed-off and replaced with fresh medium for further incubation until sample collection (1h for ChIP, 6-24 h for RNA).

### qPCR

RNA was isolated using Zymo MiniPrep RNA prep kit (R1055, Zymo Research) with DNase I treatment. The total RNA was reverse-transcribed using SuperScript III Reverse Transcriptase (Life Technologies, 18080-051) with random hexamer and oligo-dT primers as per the manufacturer’s instructions. qPCR was performed on Mx3000P qPCR systems (Agilent) using VeriQuest FastSYBR 2X qPCR master mix (Thermo, C-75690). Normalization of expression was done using GAPDH or ACTB mRNA as internal controls. For all RT-qPCR and ChIP-qPCR, experiments were performed with at least two replicates, and technical duplicates or triplicates for each biological sample. A list of primers used for qPCR is provided. If not indicated otherwise, p values were obtained using a two-tailed Student’s t-test, * = p<0.05 and ** = p<0.01, or as indicated.

### Western Blot

Cells were harvested in 350 µl of RIPA buffer (Thermo, 89900) supplemented with Protease and phosphatase inhibitor cocktail (Thermo, 78440) and tip-sonicated (Branson sonifier, 10 s) on ice. The lysate was centrifuged at 10000 rpm for 10 min at 4°C and the supernatants were used fresh or stored at -80Western blotting was performed as described(Brandl, Wagner et al. 2012) with modifications using Xcell II™ Electrophoresis and Blot System (Thermo), using precast Bis-Tris Gels (NP0323BOX, Thermo) and MOPS SDS Running buffer (MB1035, Biopioneer) for SDS-PAGE followed by transfer to Nitrocellulose membranes (1620115, BioRad) in Transfer buffer (MB1037, Biopioneer). Membranes were blocked in 5% BSA or non-fat dry milk in PBS + 0.05% Tween-20, incubated with primary antibodies 4 h to over-night at 4°C and with HRP-coupled secondary antibodies 1h at room temperature in 2% BSA or NFDM in PBS-Tween. ECL substrates (34580, Thermo) were used for signal detection with X-ray film (MUBF-02, Biopioneer) with an automated film developer (Bio pioneers) or a chemiluminescent imaging system (Pxi, Syngene).

### Chromatin Immunoprecipitation (ChIP) and ChIP-seq

ChIP was performed as previously described ref(Oh, Shao et al. 2021). In brief, cells were either cross-linked directly with 1% formaldehyde at room temperature for 10 mins. The cross-linking was quenched by the addition of 0.125M glycine for 10 min. Chromatin was fragmented using sonication (Bioruptor Pico, Diagenode) (10–30 cycles, 30 s on/30 s off). Subsequently, the soluble chromatin was cleared by centrifugation (10000 g, 10 min 4°C), pre-cleared with 10-20 μl Protein G Dynabeads (10009D, Life Technologies), and then incubated with 2 μg of antibodies at 4 °C overnight. ChIP complexes were collected using 30 μl of Protein G Dynabeads per sample for the last hour of incubation. The immune complexes were subjected to washes once with wash buffer I, twice with wash buffer II, once with Tris-EDTA (TE) + 0.1% Triton X-100, and once with TE, and then the beads were incubated at 55 °C for 2 h with proteinase K and de-cross-linked at 65 °C overnight. The final ChIP DNA was extracted and purified using QIAquick columns. For ChIP-seq, the extracted DNA was ligated to specific adaptors for Illumina’s system using the KAPA Hyper Prep Kit (Kapa Biosystems).

### RNA-seq

RNA-seq data were generated as previously described(Godoy, Niesman et al. 2019). In brief, total RNA either UT or H_2_O_2_ treated cells was extracted using the Rneasy mini RNA-isolation kit. The quality was assessed based on RNA integrity number (RIN) using an Agilent Bioanalyzer. Libraries were prepared using the Illumina TruSeq stranded mRNA kits and sequenced using Illumina NovaSeq 6000. Samples were sequenced to an average of around 25 million read pairs.

## CONTRIBUTIONS

The study was conceived and initiated by PP and MGR; experiments presented in the manuscript were performed by PP. Mouse colonies were maintained by HT RNA-seq Library prep was performed by KO. AG performed bioinformatics analysis. PP and MGR wrote the manuscript.

## ACKNOWLEDGEMENTS

We thank the UCSD mouse facility, Kristen Jepsen and the UCSD IGM core for sequencing. We thank Janet Hightower for helping with figure preparation. This publication includes data generated at the UC San Diego IGM Genomics Centre utilising an Illumina NovaSeq 6000 that was purchased with funding from a National Institutes of Health SIG grant (#S10 OD026929). This study was supported by the Larry L. Hillblom Foundation to PP (#67779) and by National Institute of Diabetes, Digestive, and Kidney Diseases and the National Heart, Lung, and Blood Institute R01HL150521 MGR.

---

**Supplementary Figure 1.**
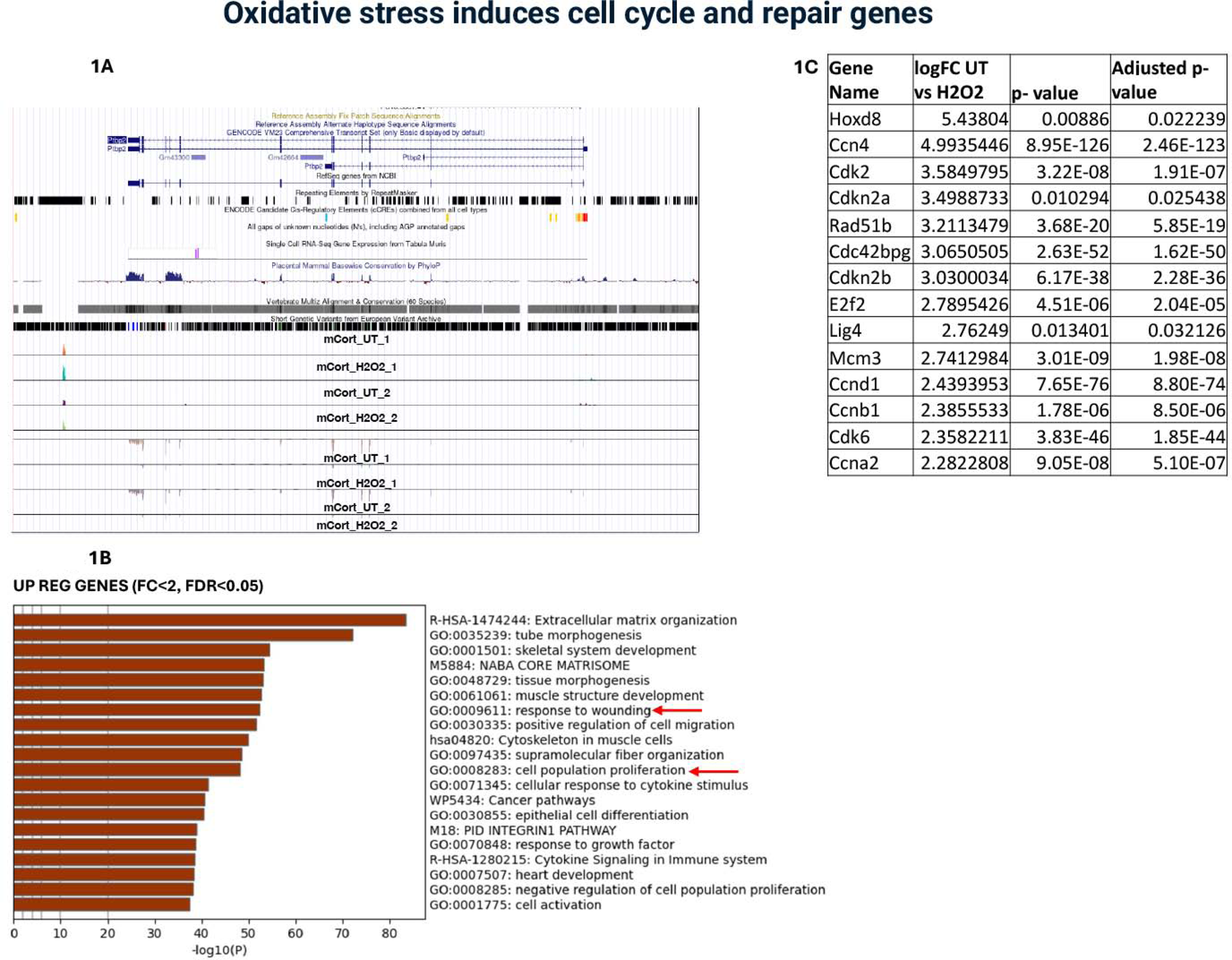
Oxidative stress induces cell cycle and repair-associated gene expression. (A–C) Transcriptomic and expression analyses of genes upregulated under oxidative stress conditions (fold change <2, FDR>0.05), GO term analysis showing enrichment of cell cycle and DNA repair pathways. Genome browser tracks (mCort_UT vs mCort_H_2_O_2_) illustrate increased transcriptional expression of PTBP1 under H_2_O_2_ stress conditions.

**Supplementary Figure 2.**
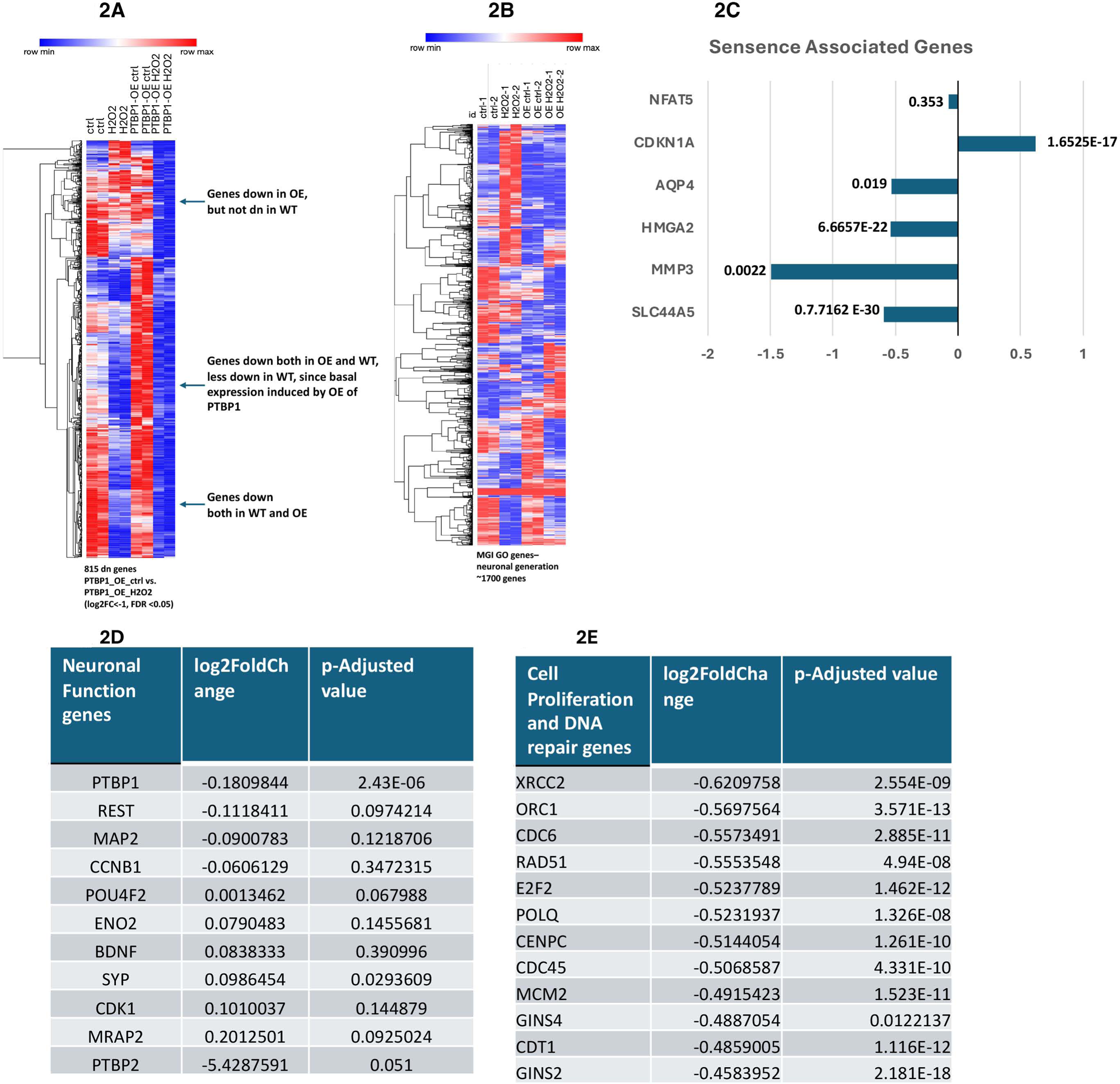
PTBP2 depletion does not recapitulate PTBP1-mediated transcriptional reprogramming. (A) RNA-seq analysis heatmap showing the comparisons of PTBP1_OE_control cells vs the PTBP1_OE_H_2_O_2_, which identified 815 downregulated genes. This shows a subset that was either downregulated in PTBP1_OE cells but not in control cells. (B) RNA-seq analysis heatmap of gene expression of a total of ∼1700 genes belonging to neuronal generation pathways as per the MGI, showing that PTBP1 OE drastically changes this subset of neuronal genes. (C-E) Relative Log2 Fold change expression of genes associated with senescence-associated genes plotted from RNA-seq analysis in either control siRNA or siRNA targeting PTBP2 in mouse primary cortical neurons. Neuronal function and stress-responsive genes following PTBP2 depletion, demonstrating that PTBP2 knockdown does not phenocopy the transcriptional effects observed upon PTBP1 overexpression.

## Notes

### Competing Interest Statement

The authors have declared no competing interest.

